# Determining the antiviral mechanism of MARCH2

**DOI:** 10.1101/2023.09.18.558306

**Authors:** Supawadee Umthong, Uddhav Timilsina, Mary D’Angelo, Spyridon Stavrou

## Abstract

Membrane-associated RING-CH (MARCH) 2 protein is a member of the MARCH protein family of RING-CH finger E3 ubiquitin ligases that have important functions in regulating the levels of proteins found on the cell surface. MARCH1, 2 and 8 inhibit HIV-1 infection by preventing the incorporation of the envelope glycoproteins in nascent virions. However, a better understanding on the mechanism utilized by MARCH proteins to restrict HIV-1 is needed. In this report, we identify an amino acid in human MARCH2, that is absent in mouse MARCH2, critical for its antiretroviral function. Moreover, we map the domains of human MARCH2 critical for restricting as well as binding to the HIV-1 envelope glycoproteins. Our findings reveal important new aspects of the antiviral mechanism utilized by human MARCH2 to restrict HIV-1 that have potential implications to all MARCH proteins with antiviral functions.

## Introduction

Retroviruses are a diverse family of viruses that infect numerous species and have the unique ability of integrating inside the host’s genome. Therefore, mammalian cells have developed defense mechanisms against them including proteins that can counteract every step of the retrovirus life cycle (1, 2). Among these host factors are the Membrane associated RING-CH (MARCH) family proteins, which are E3 ubiquitin ligases with important cellular functions (3, 4). MARCH proteins are highly conserved, are found in all mammals, with mice and humans having 11 MARCH family members (5, 6) and have critical immunomodulatory functions by regulating the levels of various immune receptors (e.g., CD86/B7.2, MHC-II etc.) found on the cell surface (3, 4).

The functions of MARCH protein family members extend beyond their role as immune receptor modulators. MARCH proteins have emerged as important antiviral factors that target a number of enveloped viruses (7). Enveloped viruses contain a lipid bilayer on the surface of the viral particle, in which viral proteins critical for virus entry localize. In the case of Human Immunodeficiency Virus type 1 (HIV-1), the viral proteins found on the surface of the cell form a heterotrimer consisting of the surface (SU) glycoprotein gp120 and the transmembrane (TM) glycoprotein gp41 (8). MARCH1, 2 and 8 exert their antiviral effect by preventing envelope glycoprotein incorporation into nascent virions for a number of viruses including HIV-1, Vesicular Stomatitis Virus (VSV), SARS-CoV-2, Influenza A virus and others, (9–13). Nevertheless, the mechanism of restriction has not been elucidated and seems to be specific to the envelope glycoproteins under investigation (10, 11, 14, 15). Finally, MARCH8 is highly expressed only in monocyte derived macrophages (MDMs), a cell type infected by HIV-1, and can inhibit HIV-1 infection at endogenous levels (9). Thus, MARCH8 is an important factor in the cellular defense against HIV-1 infection in terminally differentiated myeloid cells.

While a number of studies have investigated the antiviral role of MARCH8, less is known about the role of the other MARCH proteins vis à vis retrovirus restriction. MARCH1 and 8 form a subgroup, because of the high degree of sequence and structural homology, while MARCH2 is more similar to MARCH3, a MARCH protein with no antiretroviral function (4, 15, 16). We previously compared the human and murine orthologs of MARCH1, 2 and 8 and found that while mouse MARCH2 is about 95% homologous to human MARCH2, it has no antiviral function against retroviruses including HIV-1 and murine leukemia virus (MLV) (10), suggesting that MARCH2 acquired its antiviral function later on in the evolutionary scale.

In this report, we investigated the importance of MARCH2 during retrovirus infection. We demonstrate that Gly61 found in human MARCH2, but is absent in mouse MARCH2, is critical for its anti-HIV-1 effect. Furthermore, we determine the MARCH2 domains critical for restriction and map the interaction between MARCH2 and the HIV-1 envelope (Env).

## Results

### Transcriptional regulation of *MARCH2*

Murine *March2* expression levels are not affected by type I interferon (IFN) stimulation or MLV infection (10). In contrast, a previous report showed that human *MARCH2* mRNA levels are modestly upregulated by IFNα, a type I IFN, in MDMs (15). Thus, we examined the effect of type I IFN on *MARCH2* expression levels in cell lines susceptible to HIV-1 infection, H9 (ATCC), a human T lymphocytic cell line, and phorbol 12-myristate 13-acetate (PMA)-differentiated THP-1 (ATCC), a human monocytic cell line, after treatment with human IFN-β (500 U/ml) (PBL Assay Science). We also included 293T cells, a cell line routinely used in HIV-1 research. We found that *MARCH2* transcript levels were unaffected upon IFN-β treatment in all cell lines tested (Fig 1a). As a positive control, we used *ISG15*, an IFN-stimulated gene (17) (Fig S1a). We previously showed that MLV infection, did not affect *March2* expression levels in both primary cells and established cell lines (10). Thus, we examined the effect of HIV-1 infection on the RNA levels of *MARCH2* in H9 and THP-1 cells. We infected H9 cells and PMA-differentiated THP-1 cells with an X4-tropic (HIV-1^NL4-3^) and an R5-tropic (HIV-1^JR-CSF^) HIV-1 strain respectively and isolated RNA from the infected cells at various time points to measure *MARCH2* RNA levels. To ensure that our cells are infected, we initially measured HIV-1 *env* and *nef DNA* levels at different time points (Fig S1b). We found that HIV-1 infection had no effect on *MARCH2* expression in both cell lines tested (Fig 1b). Thus, we concluded that similar to *March2*, *MARCH2* is not induced by type I IFN or HIV-1 infection.

**Fig 1.**
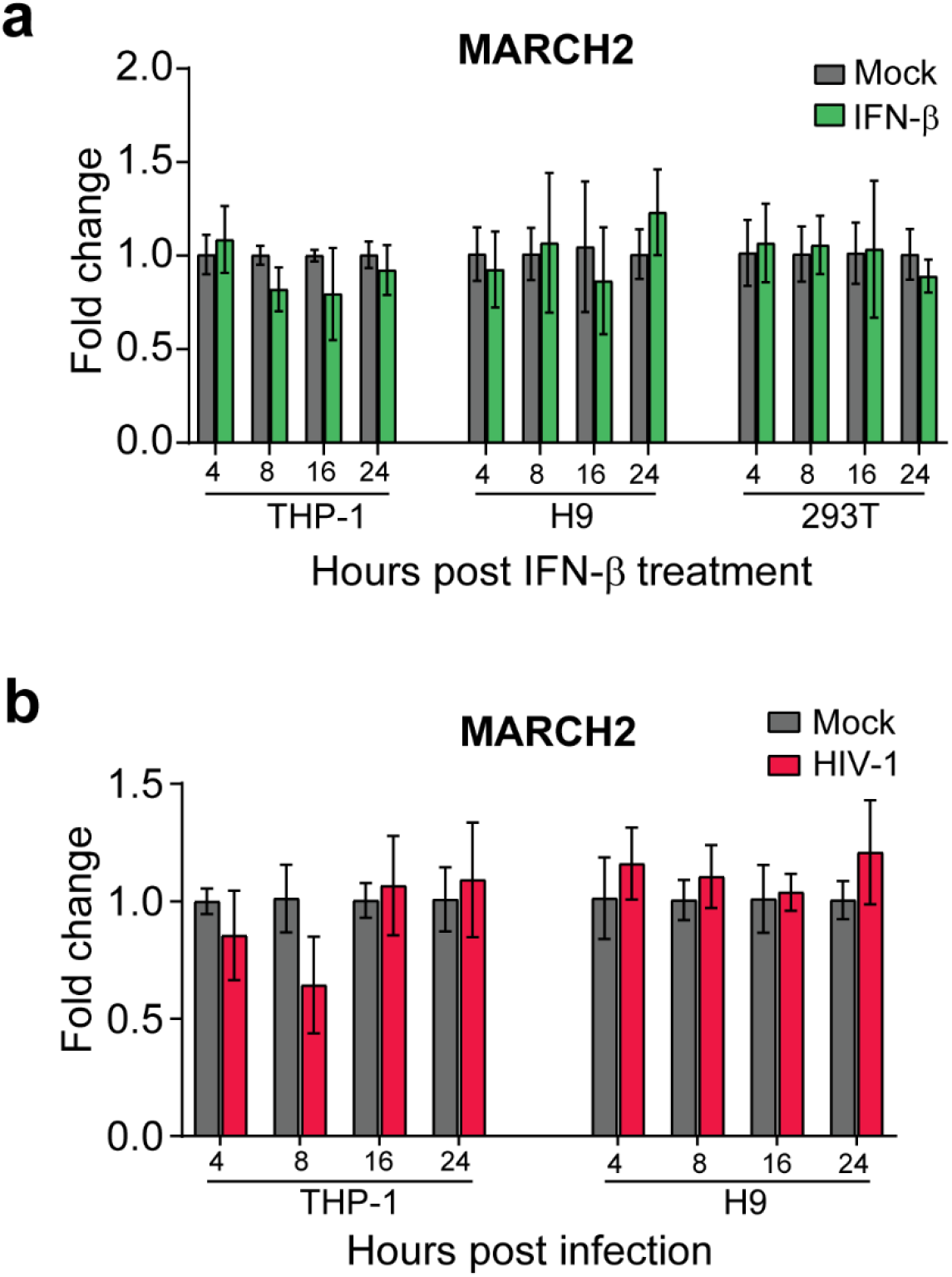
Human *MARCH2* gene expression is unaffected by type I IFN and HIV-1 infection. **(a)** Fold expression change of human *MARCH2* transcript relative to mock, normalized to GAPDH in PMA-differentiated THP-1, H9, and 293T cells treated with human IFN-β (500 units/ml) for 4 h, 8 h, 16 h, and 24 h. **(b)** Human *MARCH2* expression fold change relative to mock, normalized to GAPDH in PMA-differentiated THP-1 cells infected with HIV-1^JR-CSF^ and H9 cells infected with HIV-1^NL4-3^ at 4 h, 8 h, 16 h, and 24 h postinfection. Mock indicates mock-treated (PBS). All graphs represent mean ± SD from 3 independent experiments.

### The N-terminal cytoplasmic tail of human MARCH2 is critical for its antiviral function

We previously showed that human MARCH2 inhibits HIV-1 infection, while mouse MARCH2 has no effect (10). To investigate whether the difference in restriction phenotype could be clarified on the basis of genetic differences between *MARCH2* genes from different species, we initially compared the amino acid sequences of the human and murine MARCH2. When we compared the human and mouse MARCH2 protein sequences, we observed that these orthologs shared 95% amino acid sequence identity with all sequence differences clustered within their N- and C-terminal cytoplasmic tails (Fig 2a). To determine which residues of human MARCH2 are important for its antiretroviral phenotype, we initially made a number of MARCH2 chimeras, in which we swapped the N-terminal half of human MARCH2 with that of mouse MARCH2 and vice versa (mMARCH2^N^–hMARCH2^C^ and hMARCH2^N^–mMARACH2^C^) (Fig S2a). Cells were co-transfected with either empty vector (E.V.), or plasmids expressing human MARCH2, mouse MARCH2 or the aforementioned MARCH2 chimeras along with either an MLV or HIV-1 molecular clone (NL4-3). Cells and media were harvested 48 hours post-transfection followed by western blots to determine the levels of the MLV (gp70/SU, p15E/p13E/TM) and HIV-1 (gp120/SU, gp41/TM) envelope glycoproteins. In the cell fractions, we found that the chimera containing the N-terminal end of human MARCH2 (hMARCH2^N^–mMarch2^C^) resulted in the reduction of both MLV and HIV-1 envelope glycoprotein levels similar to that seen with human MARCH2. On the other hand, the chimera containing the N-terminal end of mouse MARCH2 (mMarch2^N^–hMARCH2^C^) had no effect on the viral envelope glycoprotein levels, similar to mouse MARCH2 (Fig 2b, c). Similarly, MLV and HIV-1 virions produced in the presence of either human MARCH2 or hMARCH2^N^–mMarch2^C^ had significantly lower levels of gp70/p15E and gp120/gp41 respectively compared to virions produced in the presence of either E.V., mouse MARCH2 or mMarch2^N^–hMARCH2^C^ (Fig 2b, c). Therefore, we concluded that the N-terminal end of human MARCH2 is critical for its antiviral effect.

**Fig 2.**
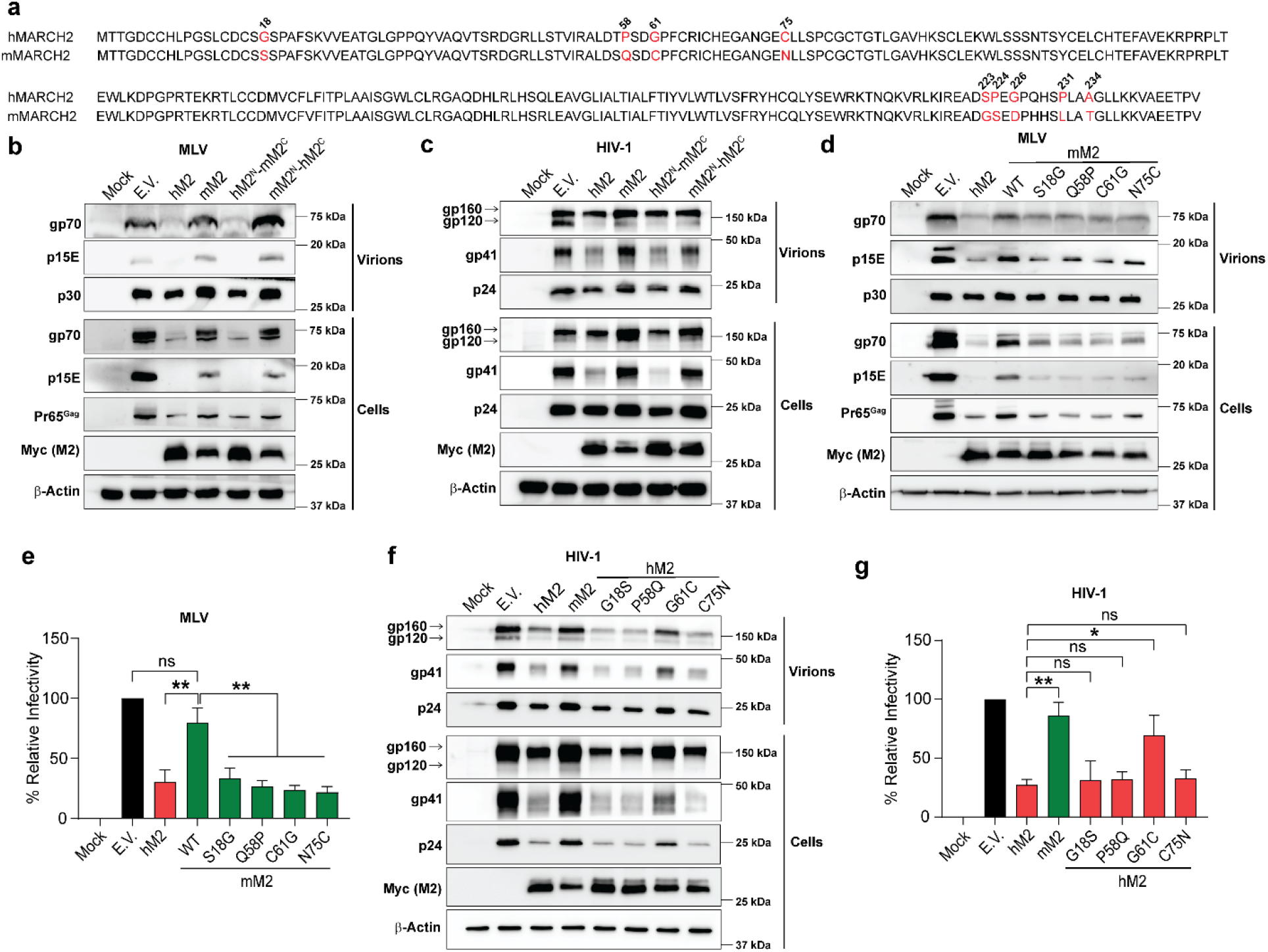
G61 residue on the N-terminal cytoplasmic tail of human MARCH2 is critical for its antiretroviral function. **(a)** Sequence alignment of human MARCH2 (hMARCH2: NCBI Ref. NP_001356705.1) and mouse MARCH2 (mMARCH2: NCBI Ref. NP_001239409.1. Differing amino acid residues are shown in red and positions are indicated. **(b and c)** The N-terminal cytoplasmic tail of human MARCH2 is essential for its antiretroviral function. 293T cells were co-transfected with molecular clones for MLV (FMLV) (**b**) or HIV-1 (NL4-3) (**c**) along with either wildtype MARCH2 (hM2 or mM2), MARCH2 chimeras, in which the cytoplasmic tails have been swamped swapped (hM2^N^-mM2^C^ or mM2^N^-hM2^C^), or empty vector (E.V.). At 48 h post-transfection, cells and released virus in the culture medium were harvested and analyzed by immunoblotting. **(d and e)** Substituting any 4 N-terminal cytoplasmic tail residues (S18, Q58, C61, and N75) of mouse MARCH2 with that from human MARCH2 renders mouse MARCH2 antiviral. In **(d)**, 293T cells were co-transfected with an infectious clone of MLV along with either wildtype human or mouse MARCH2 along,the indicated mouse MARCH2 mutants (S18G, Q58P, C61G, or N75C) and E.V. Cells and released virus in the culture medium were analyzed by immunoblotting as mentioned above. In **(e)**, MLV containing firefly luciferase reporter genome was generated as described in (**d**) and was used to infect NIH 3T3 cells, luciferase levels were measured 48 hpi and normalized to MLV p30^CA^ levels of the input virus. The percentage (%) of relative infectivity was determined with respect to virus produced in the presence of E.V. **(f and g)** G61 residue at the N-terminal cytoplasmic tail of human MARCH2 is critical for its anti-HIV-1 function. In (**f**), 293T cells were co-transfected with the HIV-1 molecular clone, NL4-3, along with either wildtype mouse or human MARCH2s, human MARCH2 mutants (G18S, P58Q, G61C, or C75N) or E.V. Cells and released virus in the culture medium were processed as described above and analyzed by immunoblotting. In (**g**), NL4-3 Env pseudotyped luciferase reporter viruses were generated in the presence of wildtype human and human MARCH2, human MARCH2 mutants (G18S, P58Q, G61C, or C75N) or E.V. Pseudoviruses were used to infect U373-MAGI-CXCR4 cells, luciferase levels were measured 48 hpi and normalized to HIV-1 p24^CA^ levels of the input virus. The percentage (%) of relative infectivity was determined with respect to virus produced in the presence of E.V. For **b**, **c**, **d** and **f**, representative immunoblotting results from n = 3 independent experiments are shown. All graphs represent mean ± SD from 3 independent experiments. For **e** and **g**, statistical significance was determined using one-sample t-test (two-tailed) when comparisons were performed with E.V. and unpaired t-test (two-tailed) was used for non E.V. comparisons. ns, non-significant; *, *P* ≤ 0.05; **, *P* ≤ 0.01. (human MARCH2, hM2; mouse MARCH2, mM2; wildtype, WT)

There are four residues (18, 58, 61 and 75) in the N-terminal cytoplasmic tail of human and mouse MARCH2 that are radically different between the two orthologs and thus may cause significant changes in the MARCH2 protein structure and function (Fig 2a). While residue 18 is located outside of the RING-CH domain, the other three (58, 61, and 75) are positioned close to it (Fig S2a). To determine which of these residues is/are important for the antiretroviral effect of human MARCH2, we made a number of human and mouse MARCH2 mutants, in which we replaced, by site-directed mutagenesis, the amino acids found at the aforementioned sites in mouse MARCH2 to those found in human MARCH2 (S18G, Q58P, C61G, and N75C) and vice versa in human MARCH2 (G18S, P58Q, G61C, and C75N). Initially, we co-transfected 293T cells with an MLV infectious clone and either human MARCH2, mouse MARCH2, the mouse MARCH2 mutants mentioned above (mMARCH2 S18G, mMARCH2 Q58P, mMARCH2 C61G, and mMARCH2 N75C) or E.V. We found that all mouse MARCH2 mutants tested reduced gp70 and p15E levels in the cellular and viral fractions similar to the levels seen with human MARCH2 (Fig 2d). In agreement with our western blot data, all mouse MARCH2 mutants reduced MLV particle infectivity similar to that observed with human MARCH2 (Fig 2e). Therefore, we concluded that substituting any of these four sites in mouse MARCH2 with the amino acids found in human MARCH2 is sufficient to render mouse MARCH2 antiviral against MLV. To determine the residue of human MARCH2 critical for its anti-HIV-1 function, we transfected cells with the NL4-3 molecular clone and the human MARCH2 mutants carrying the mouse MARCH2 amino acids at the same sites (hMARCH2 G18S, hMARCH2 P58Q, hMARCH2 G61C, and hMARCH2 C75N) or E.V. We found that when human MARCH2 carried the amino acid of the mouse ortholog at position 61 (hMARCH2 G61C), it lost its antiviral effect against HIV-1 (Fig 2f). We also measured HIV-1 particle infectivity in the presence of the different human MARCH2 mutants and found, in agreement with our western blot data, that human MARCH2 G61C had lost its ability to inhibit viral particle infectivity similar to mouse MARCH2 (Fig 2g). Knowing the critical role of amino acid residue at position 61 of human MARCH2 and the anti-MLV activity of the mouse MARCH2 C16G mutant, we then asked if mouse MARCH2 C16G mutant can also inhibit HIV-1. For this we measured infectivity of HIV-1 particles produced in the presence of mouse MARCH2 C61G mutant and found that unlike what was observed with MLV, C61G substitution in mouse MARCH2, similar to what we found with mouse MARCH2, had no effect on HIV-1 particle infectivity (Fig. S2b). In summary, our findings demonstrate that Gly61 at the N-terminal cytoplasmic tail of human MARCH2 is critical for its anti-HIV-1 function. On the other hand, substituting any of the four residues in mouse MARCH2 N-terminal cytoplasmic tail to the amino acids found in human MARCH2 rendered mouse MARCH2 antiviral towards MLV.

### Naturally occurring MARCH2 polymorphisms potently restrict HIV-1 envelope

Single nucleotide variants (SNVs) in cellular restriction factors can dramatically influence their ability to counteract HIV-1 infection. Previous reports have identified SNVs in TRIM5α and APOBEC3D that affect their anti-HIV-1 function (18–20). To identify SNVs in the *MARCH2* locus, we accessed the 1000 Genome Project Phase III genotypes for the coding region of the *MARCH2* gene (n=5008). We investigated for missense SNVs that occur at high frequency and would lead to radical changes in the amino acid sequence of MARCH2 protein. Two SNVs that fit our search criteria were an alanine changed to a threonine at position 54 (A54T/rs1133893) and an arginine changed to a proline at position 219 (R219P/rs34099346). The minor allele frequency of A54T was the highest in European and American populations (0.2922 and 0.272 respectively) and that of R219P was the highest in South Asian populations (0.174) (Supplemental Table S1). To determine the effect of these SNVs on the antiviral function of MARCH2, we generated NL4-3 Env pseudotyped luciferase reporter HIV-1 viruses in the presence of varying amounts of wildtype MARCH2 or MARCH2 carrying the two SNVs (MARCH2 A54T and MARCH2 R219P). At first, we examined the effect of these MARCH2 SNVs on the incorporation of HIV-1 Env glycoproteins in nascent virions by performing western blots on purified viral pellets. Interestingly, we found that compared to virions produced in the presence of wildtype MARCH2, virions generated with MARCH2 carrying the two SNVs had significantly lower gp120 and gp41 levels at all concentrations tested (Fig S3a). Next, we examined the infectivity of these pseudoviruses by infecting U373-MAGI-CXCR4 cells and found that, in agreement with our western blot findings, MARCH2 carrying the two SNVs reduced HIV-1 particle infectivity more potently than wildtype MARCH2 (Fig S3b). Thus, these two polymorphisms (A54T/rs1133893 and R219P/rs34099346) impart gain-of-function on MARCH2 regarding its antiviral effect.

### Mapping the domains of MARCH2 responsible for its anti-HIV-1 function

MARCH2 has two cytoplasmic tails, an N-terminal tail, in which the RING-CH domain is located, and a C-terminal tail, which contains the PDZ binding motif, along with two transmembrane (TM) domains (5, 21). To investigate the role of these domains in MARCH2-mediated anti-HIV-1 activity, we generated a series of constructs with mutations in the aforementioned domains. Initially, we confirmed that the mutations we introduced in MARCH2 did not affect its localization with cellular membranes by performing western blots on lysates of membrane bound proteins isolated from 293T transfected with either the wildtype or mutant MARCH2 expression plasmids, as antibodies available for MARCH2 are not suitable for flow cytometry. We found that all MARCH2 mutants tested, localized to cellular membranes resembling wildtype MARCH2 (Fig S4a). To examine the role of the RING-CH domain on the anti-HIV-1 activity of MARCH2, we generated a MARCH2 mutant by introducing a mutation (W97A), which abrogates the activity of the RING-CH domain (15). We observed that in the presence of MARCH2 W97A, the levels of gp120 and gp41 were similar to those seen in the presence of E.V. (Fig 3a). When using MARCH2 constructs with deletions at the N-terminal cytoplasmic tail upstream of the RING-CH domain (Δ2-30 and Δ31-56 amino acids), we noticed that MARCH2Δ2-30 and MARCH2Δ31-56 reduced gp120 and gp41 levels similar to wildtype MARCH2 (Fig 3b). Likewise, mutations in the PDZ binding motif (243-ETVA-246 to 243-AAAA-246) did not affect the ability of MARCH2 to reduce gp120 and gp41 levels (Fig 3c). To investigate the role of the MARCH2 TM domains on retrovirus restriction, we generated a series of mutants, in which we substituted the TM domains of MARCH2 with those of MARCH4 that has no anti-HIV-1 activity (15). Interestingly, we found that the exchange of the second TM domain of MARCH2 with that from MARCH4 affected protein stability (data not shown). Therefore, for the second TM domain of MARCH2, we replaced it with the TM domain of the human transferin receptor (TR). Upon verification that swapping the TM domains of MARCH2 did not affect the localization of MARCH2 to cellular membranes (Fig S4b), we found that mutating just the second TM domain of MARCH2 rendered it unable to reduce the gp120 and gp41 levels (Fig 3d). The importance of the second TM domain is further supported by our studies on the two isoforms of MARCH2, which are products of alternative splicing. The canonical long isoform of MARCH2 (*March2-001*) has two TM domains, while the non-canonical shorter isoform (*March2-002*) lacks both TM domains (Δ125-194) (Fig S5). We initially verified the expression of both isoforms in H9 and THP-1 cells by PCR (Fig 4a). Subsequently, we generated NL4-3 Env pseudotyped luciferase reporter viruses in the presence of either the long or the short MARCH2 isoform and examined particle infectivity by infecting U373-MAGI-CXCR4 cells. We found that in the presence of the long MARCH2 isoform viral particle infectivity was reduced, while the shorter MARCH2 isoform had no effect (Fig 4b). In conclusion, our data show that the second TM domain of MARCH2 is critical for its antiviral function.

**Fig 3.**
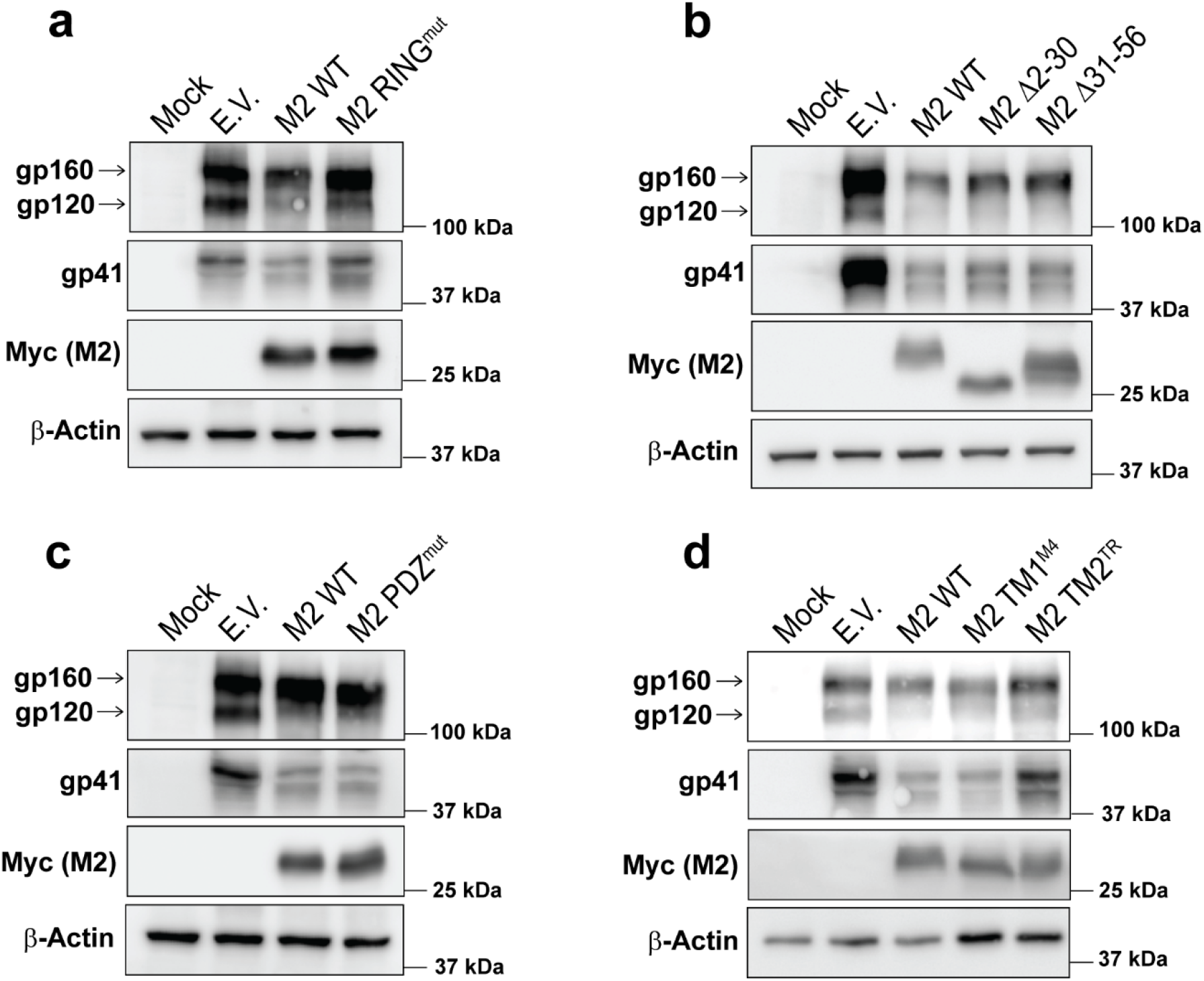
The RING-CH and the TM2 domain of human MARCH2 are important for the MARCH2 antiviral effect. 293T cells were co-transfected with an HIV-1 molecular clone (NL4-3) along with either empty vector (E.V.), wildtype human MARCH2 (M2 WT) or mutant human MARCH2. **(a)** MARCH2 with a mutated RING-CH domain (M2 RING^mut^), **(b)** MARCH2 with deletions in N-terminal tail (M2 Δ2-30; M2 Δ31-56), **(c)** MARCH2 with mutations in the PDZ binding motif (M2 PDZ^mut^) and **(d)** MARCH2 with either N-terminal (TM1) or C-terminal (TM2) swapped with those of human MARCH4 (M2 TM1^M4^) and human transferrin receptor (M2 TM1^TR^). Cells were harvested 48 h post-transfection, and proteins were analyzed by immunoblotting. For **a-d**, representative immunoblotting results from n = 3 independent experiments are shown.

**Fig 4.**
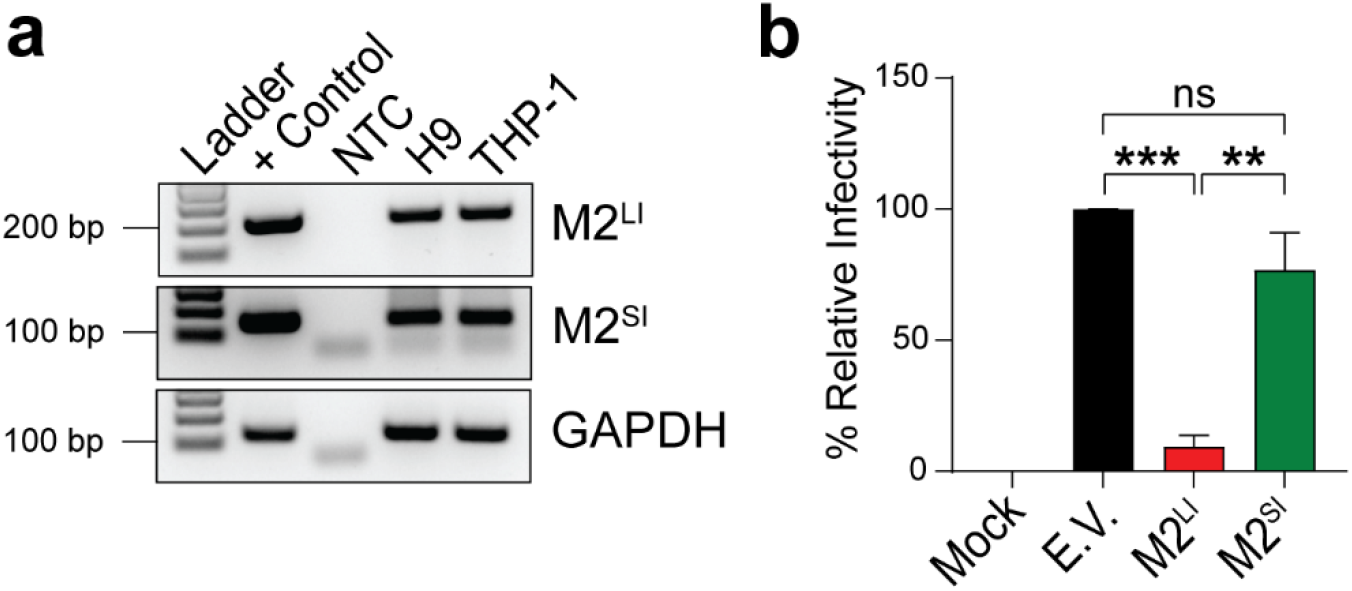
The canonical long isoform of MARCH2 has anti-HIV-1 function. **(a)** Both isoforms of human MARCH2 are expressed in H9 and THP-1 cells. H9 and THP-1 cDNA was used to detect the two MARCH2 isoforms. DNA from MARCH2 transfected cells served as positive control. **(b)** Canonical long isoform of human MARCH2 reduces HIV-1 infectivity. NL4-3 Env pseudotyped luciferase reporter viruses were generated in the presence of either the canonical long isoform of MARCH2 (M2^LI^), the non-canonical shorter isoform (M2^SI^) or empty vector (E.V.). Pseudoviruses were used to infect U373-MAGI-CXCR4 cells, luciferase levels were measured 48 hpi and normalized to HIV-1 p24^CA^ levels of the input virus. The percentage (%) of relative infectivity was determined with respect to virus produced in the presence of E.V. In **a**, representative gel images from *n* = 2 independent experiments are shown. In **b**, graphs represent mean ± SD from 3 independent experiments. Statistical significance was determined using one-sample t-test (two-tailed) when comparisons were performed with E.V. and unpaired t-test (two-tailed) was used for comparison between M2^LI^ and M2^SI^. ns, non-significant; **, *P* ≤ 0.01; ***, *P* ≤ 0.001. (no template control, NTC; longer isoform of MACRH2, M2^LI^; shorter isoform of MACRH2, M2^SI^; + Control, positive control)

### MARCH2 interacts with HIV-1 gp41

MARCH proteins form complexes with their cellular or viral targets (10, 12, 22, 23). However, all previous studies have been focused on MARCH8 and only in the context of overexpression (9, 10, 12). Therefore, we set out to determine if HIV-1 Env interacts with endogenously expressed MARCH2 in immune cells. Consequently, we performed coimmunoprecipitations (coIPs) using cell lysates from HIV-1 infected H9 cells, which express endogenous levels of MARCH2. We found that endogenous MARCH2 coIPed with HIV-1 gp41 during infection (Fig 5a). To determine whether MARCH2 interacts with HIV-1 gp41 via the TM domains, we co-transfected 293T cells with an NL4-3 molecular clone and either E.V., wildtype MARCH2 or the aforementioned TM mutant MARCH2 constructs (Fig 3d). To ensure that MARCH2 did not dramatically reduce the HIV-1 envelope glycoprotein levels impacting our coIP studies, we used lower concentrations of the MARCH2 expression plasmids. We then performed coIPs pulling down for either HIV-1 gp41 or MARCH2. We found that only wildtype MARCH2 and MARCH2 that had the first TM domain mutated interacted with HIV-1 gp41 (Fig 5b). Swapping the second TM domain of MARCH2 to that of human TFR abolished that interaction (Fig 5b). Thus, our data show that MARCH2 interacts with the HIV-1 envelope glycoproteins via the second TM domain. To better understand the interaction between HIV-1 gp41 and MARCH2, we investigated the role of the TM domain of gp41 by generating a mutant HIV-1 envelope expressing plasmid, in which the gp41 TM domain is replaced with that of human TR (NL4-3 Env TM^TR^). To ensure that mutating the TM domain of HIV-1 gp41 did not affect gp41 localization at the plasma membrane, we transfected 293T cells with either the wildtype (NL4-3 Env) or the TM mutant HIV-1 envelope (NL4-3 Env TM^TR^) followed by flow cytometry using an anti-HIV-1 antibody (VRCO1) (24). We observed similar cell surface expression levels for both NL4-3 Env and NL4-3 Env TM^TR^ (Fig S6) indicating that mutating the TM domain of HIV-1 gp41 did not alter gp41 plasma membrane localization. We then examined whether NL4-3 Env TM^TR^ is resistant to MARCH2 restriction by co-transfecting 293T cells with either NL4-3 Env or NL4-3 Env TM^TR^ along with MARCH2 followed by western blots. We found that NL4-3 Env TM^TR^ was resistant to MARCH2-mediated inhibition (Fig 5c). We then performed coIPs using cell lysates from 293T cells co-transfected with either NL4-3 Env or NL4-3 Env TM^TR^ expression plasmids along with low concentration of MARCH2. We noticed that mutating the TM domain of HIV-1 gp41 abolished the interaction between MARCH2 and gp41 (Fig 5d). Taken together, our findings show that endogenous MARCH2 and HIV-1 gp41 interact and their association is mediated by their TM domains.

**Fig 5.**
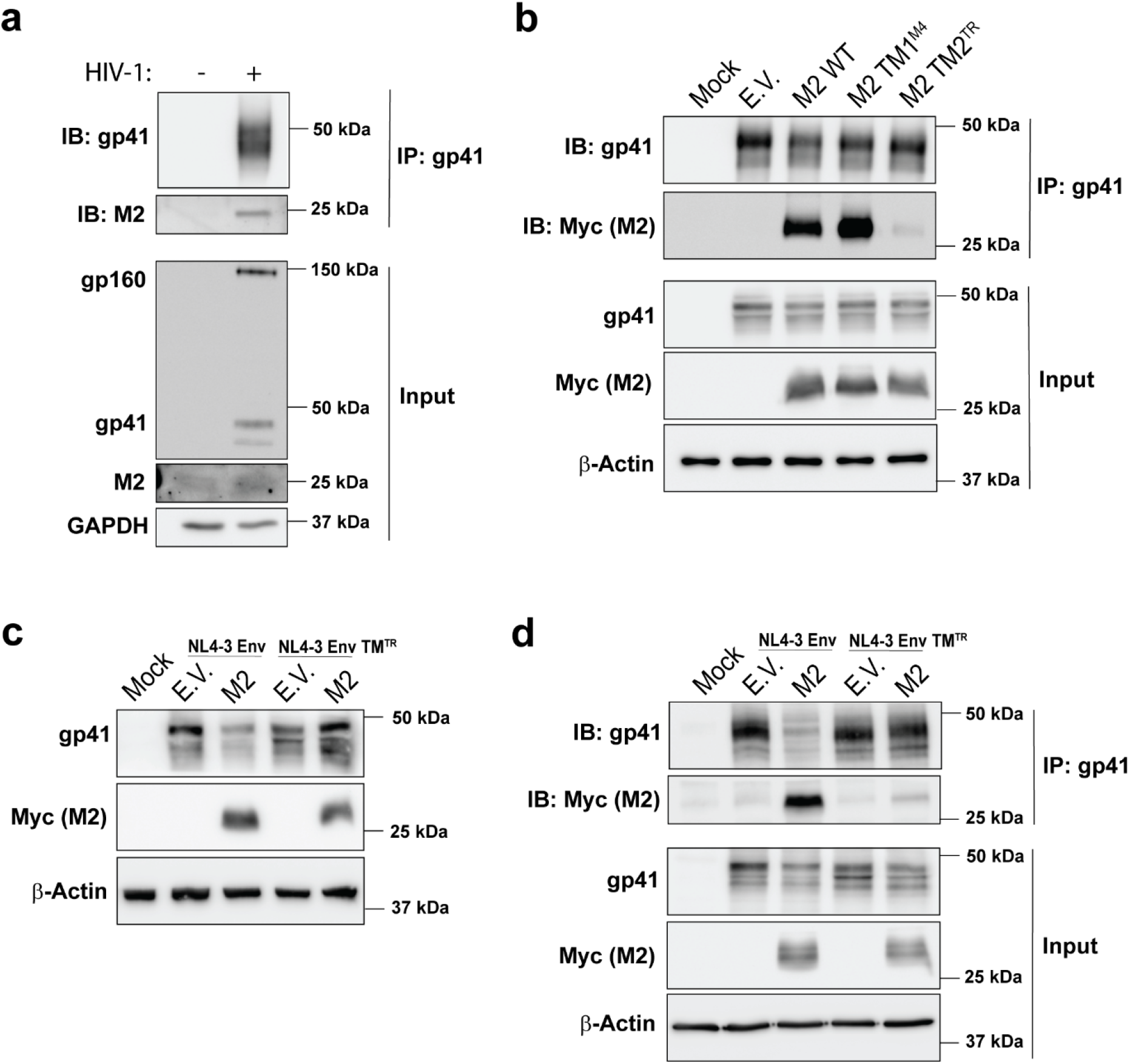
MARCH2 and HIV-1 gp41 interaction is mediated by their TM domains. **(a)** Endogenous MARCH2 interacts with HIV-1 gp41. Cell lysates from HIV-1^NL4-3^ infected H9 cells were immunoprecipitated with anti-HIV-1 gp41 antibody followed by immunoblot analysis probing with anti-MARCH2 (endogenous MARCH2) and anti-HIV-1 gp41 antibodies. **(b)** The C-terminal TM domain (TM2) of MARCH2 is critical for its interaction with HIV-1 gp41. 293T cells were co-transfected with plasmids expressing an HIV-1 molecular clone (NL4-3) along with either empty vector (E.V.), wildtype MARCH2 (M2 WT) or MARCH2 TM domain mutants (M2 TM1^M4^, M2 TM2^TR^). Cells were harvested 48h post-transfection and lysates were immunoprecipitated with anti-HIV-1 gp41 antibody followed by immunoblot analyses probing with anti-Myc (MARCH2), and anti-HIV-1 gp41 antibodies. **(c)** HIV-1 gp41 TM domain is required for MARCH2-mediated restriction. 293T cells were co-transfected with plasmids for either NL4-3 Env or mutant NL4-3 Env in which the TM domain has been swapped with that from TFR (NL4-3 Env TM^TR^) along with either empty vector (E.V.) or MARCH2 (M2). Cells were harvested as above and the indicated proteins were analyzed by immunoblotting. **(d)** TM domain of HIV-1 gp41 is required for its interaction with MARCH2. 293T cells were co-transfected with plasmids as in (c) followed by immunoprecipitation with anti-HIV-1 gp41 antibody and immunoblot analyses probing with anti-Myc (MARCH2) and anti-HIV-1 gp41. For **a-d,** GAPDH and β-actin served as our loading control for our input blots, representative immunoblotting results from *n* = 3 independent experiments are shown. (MARCH2, M2)

## Discussion

MARCH proteins have emerged as an important family of host factors that inhibit HIV-1 infection by preventing the incorporation of viral envelope glycoproteins into nascent virions (9–11, 15). Nevertheless, many questions remain regarding the mechanism they utilize to block HIV-1 infection (9, 10, 15, 25). In this study, we examine in depth the antiviral effect of MARCH2 on HIV-1 infection. MARCH2 is found ubiquitously in the host and localizes in various cellular organelles including the plasma membrane and Golgi (21, 26). Many host receptors have been identified as targets of MARCH2 including β2 andrenegic receptor and the cystic fibrosis transmembrane conductance regulator (CFTR) (27, 28). In contrast to previous findings (25), *MARCH2* RNA levels in all cell types we tested (H9 and THP-1) were unaffected by HIV-1 infection and IFN-β treatment. We speculate that the discrepancy between our findings and previous results regarding the transcriptional induction of *MARCH2* may be due to the different cell lines used. Nevertheless, in support of our data, there has been no study up to date to have reported *MARCH2* to be an ISG or HIV-1 induced gene.

Unlike MARCH 1 and 8, for which their antiviral effect is conserved in mammals, as both the murine and human homologues of MARCH1 and 8 restrict retroviruses, only human MARCH2 blocks HIV-1 infection, mouse MARCH2 has no antiviral effect (10). Interestingly the human and murine orthologs of MARCH2 differ in 10 amino acids clustered in the N- and C-terminal tails of the protein (Fig 2a). In this report, we showed that the N-terminal tail of human MARCH2 is critical for its antiviral function and identified amino acid Gly61, present in human MARCH2, but absent in mouse MARCH2, to bestow the anti-HIV-1 function in the human ortholog. The importance of a single amino acid in altering the antiviral function of host factors has also been demonstrated in the case of TRIM5α. While human TRIM5α has a very modest effect on HIV-1 replication, rhesus monkey TRIM5α potently restricts HIV-1 infection. Alteration of arginine 332 of human TRIM5α to proline, the residue found in rhesus monkey TRIM5α, renders human TRIM5α potently antiviral towards HIV-1 (29, 30). Our findings further support the notion that small changes in the amino acid sequence of proteins during evolution can have drastic effects in their antiviral phenotype.

MARCH2 is a transmembrane protein with two TM domains, a RING-CH domain (3) and a C-terminal PDZ-binding domain critical for protein-protein interactions (31, 32). In this report, we show that only the RING-CH domain and the second TM domain are critical for its antiviral function. Interestingly, the PDZ motif that is important for MARCH2-mediated protein-protein interactions (31) did not affect MARCH2 inhibition of HIV-1. In addition, we found that MARCH2 interacts with the HIV-1 envelope glycoproteins via TM-TM interactions. Transmembrane domain-mediated interactions of host restriction factors with viral proteins may have important benefits for the host due to the fact that transmembrane sequences, like that of gp41, can be quite stable and highly conserved (33), as mutations in transmembrane domains may result in structural changes affecting membrane localization of a protein thereby impacting its function (34). In addition, the importance of the MARCH2 TM domains on HIV-1 restriction is further evident, when we investigated the MARCH2 isoforms for antiviral function. Our studies showed that only the long isoform of MARCH2 (*March2-001*), which contains both TM domains, can restrict HIV-1. The fact that not all MARCH2 isoforms have antiviral function is similar to what has been seen with SERINC5 and Mx2, two other known host restriction factors (35, 36). Nevertheless, the function of the short MARCH2 isoform remains to be elucidated. In conclusion, our findings reveal new aspects of the antiviral function of the MARCH protein family and demonstrate its importance in host defense during HIV-1 infection.

## Materials and Methods

### Cell culture

293T cells (ATCC) and NIH 3T3 cells were cultured in Dulbecco’s Modified Eagle Media (DMEM; Gibco) with 10% (vol/vol) fetal bovine serum (FBS; Sigma), 6 mM L-glutamine (Gibco), and 100 mg/ml penicillin and streptomycin (P/S; Gibco) (complete DMEM). U373-MAGI-CXCR4 cells (NIH HIV Reagent Program, Division of AIDS, NIAID, NIH: ARP-3596) were cultured in complete DMEM (10% FBS, 100 mg/ml P/S) with 0.2 mg/ml G418 (Gibco), 0.1 mg/ml hygromycin B (Invitrogen), and 1.0 µg/ml puromycin (Research Products International). H9 cells (ATCC) were cultured in RPMI 1640 media (Gibco) with 10% FBS and 100 mg/ml P/S (complete RPMI). THP-1 cells (ATCC) were cultured in complete RPMI media supplemented with 0.05 mM β-mercaptoethanol.

### Plasmids

The MLV molecular clone (pLRB302), HIV-1 molecular clones: pNL4-3 and JR-CSF (pYK-JRCSF) used in this paper have been previously described (37–40). The pFB-*luc* construct has been previously described (10). The following reagents were obtained from the NIH AIDS Reagent program, Division of AIDS, NIAID, NIH: HIV-1 NL4-3 Infectious Molecular Clone (pNL4-3), ARP-2852, contributed by Dr. M. Martin; JR-CSF Infectious Molecular Clone (pYK-JRCSF), ARP-2708, contributed by Dr. S. Y. Chen and Dr. Yoshio Koyanagi and HIV-1 NL4-3 ΔEnv Vpr Luciferase Reporter Vector (pNL4-3.Luc.R-E-), ARP-3418, contributed by Dr. Nathaniel Landau.

Human MARCH2 construct has been previous described (25). Cloning strategies for human MARCH2 (hMARCH2; represents the longer isoform 1 unless stated otherwise), human MARCH4 and mouse MARCH2 (mMARCH2) into pcDNA3.1/myc-His A (Invitrogen) and pBJ5 vector have been previously described (10). Human MARCH2 isoform 2 was identified from https://www.uniprot.org/uniprotkb/Q9P0N8/entry#Q9P0N8-1/2 and was acquired from GenScript, (clone ID: OHu31844, Accession Version: NM_001005416.2) followed by cloning into pcDNA3.1/myc-His A vector using the same primers for the human MARCH2 isoform 1. MARCH2 chimeras containing N-terminal swaps (pBJ5 mMarch2^N^–hMARCH2^C^ or pBJ5 hMARCH2^N^–mMarch2^C^) were generated by using site-directed mutagenesis (SDM) to eliminate an A*pa*I cut site (nucleotide position 679) in hMARCH2 construct using a pair of hMARCH2A*pa*Iuncut primers listed in Supplemental Table S2, thus A*pa*I can only digest hMARCH2 at one site (nucleotide position 489). This modified hMARCH2 and mMARCH2 were then cut with *Xho*I-*Apa*I and *Apa*I-*Not*I to isolate the N-terminal and C-terminal fragments followed by reciprocal ligations to generate the indicated chimeras. MARCH2 mutant constructs containing point mutations and deletions used in this study were generated by SDM using Q5 polymerase (New England Biolabs) and primers listed in Supplemental Table S2. MARCH2 chimeras containing the TM domains from either human MARCH4 or human transferrin receptor (TR) were generated using the NEBuilder HiFi DNA assembly kit (New England Biolabs) and the primers listed in Supplemental Table S2. TM domain of TR was PCR amplified from 293T cDNA as template. For construction of pcDNA3.1 NL4-3 Env, we initially digested the pNL4-3 plasmid with EcoRI and XhoI and subcloned this fragment into pcDNA3.1/myc-His A (Invitrogen). To generate pcDNA3.1 NL4-3 Env TM^TR^, we replaced NL4-3 Env TM domain with that of TFR using the NEBuilder HiFi DNA assembly kit and the primers listed in Supplemental Table S2. All constructs generated were confirmed by DNA sequencing.

### Virus preparation

Virus stocks (HIV-1 NL4-3 and HIV-1 JR-CSF) were prepared by transfecting 293T cells (3 × 10^6^ cells) seeded in 10-cm-diameter cell culture dishes with 10 µg of plasmid DNAs mentioned above. Culture media was removed and replenished 24 h after transfection. All culture supernatants were harvested 48 h after transfection, centrifuged at 714 × g at 4 °C to eliminate cellular debris, filtered through 0.45 μm filter and treated with 10 U/ml DNase I (Roche) for 30 min at 37 °C. Aliquots of filtered culture supernatants were stored at −80 °C for future experiments. HIV-1 p24^Gag^ levels were determined by using a HIV-1 p24 ELISA kit (Xpress Biotech International).

### HIV-1 infection, interferon treatment and human *MARCH2* expression analysis

THP-1 cells were treated with 50 nM of phorbol 12-myristate 13-acetate (PMA) (MedChemExpress) for 2 days. PMA-differentiated THP-1 cells and H9 cells were seeded at a density of 5 × 10^4^/well in a 96-well plate and treated with or without 500 U/ml of human Interferon Beta 1β (PBL Assay Science). Cells were harvested at 4, 8, 16, and 24 h post treatment and RNA was isolated using RNeasy Mini kit (Qiagen). cDNA was synthesized using the SuperScript III First Strand Synthesis kit (Invitrogen) per manufacturer’s recommendation. RT-qPCR was performed using the PowerUp SYBR Green PCR master mix kit (Applied Biosystems) in a CFX384 Touch Real-Time PCR detection system (Bio-Rad). The following primers were used; *human MARCH2*: 5’-GCTGTCTGGAGAAGTGGCTT-3’/5’-CTTCAGCCACTCTGTGAGGG-3’, *GAPDH* 5ʹ-AACGGGAAGCTTGTCATCAATGGAAA-3ʹ/5ʹ-GCATCAGCAGAGGGGGCAGAG-3ʹ and *Interferon-stimulated gene 15* (*ISG15*): 5ʹ-GATCACCCAGAAGATCGGCG-3’/5’-GGATGCTCAGAGGTTCGTCG-3ʹ.

PMA-differentiated THP-1 and H9 cells seeded at a density of 5 × 10^4^ cells per well in a 96 well plate were infected with a 5 MOI of HIV-1 (HIV-1^JR-CSF^ and HIV-1^NL4-3^) respectively by spinoculation as previously described (41). Cells were harvested at 4, 8, 16, and 24 h post infection. RNA isolation, cDNA synthesis and RT-qPCR were performed as mentioned above. To quantify total HIV-1 DNA in the infected cells, DNA was extracted using a DNeasy blood and tissue kit (Qiagen) and RT-qPCR was performed as mentioned above using the following primers: NL4-3 *Env*: 5’-TAAAGTGCACTGATTTGAAGAATGAT-3’/ 5’-ATCTCTTATGCTTGTGCTGATATTG-3’, NL4-3 *Nef*: 5’-CAAGTGGTCAAAAAGTAGTGTGATT-3’/ 5’-ATACTGCTCCCACCCCATC-3’; JR-CSF *Env*: 5’-TGATAGTAGGAGGCTTGATAGGT-3’/5’-GAGGAGGGTCTGAAACGATAAG-3’, JR-CSF *Nef*: 5’-CACAAGGCTACTTCCCTGATT-3’/5’-CTCCTTCATTGGCCTCTTCTAC-3’. The relative levels of amplification were quantified for each sample from standard curves generated using known quantities of DNA standard templates and normalized to GAPDH levels.

### Transfection and Immunoblotting

All transfections were performed on 293T cells (seeded at a density of 0.5 × 10^6^ cells/well of a 6-well plate one day prior) using Lipofectamine 3000 transfection reagent (Thermo Fisher Scientific) according to the manufacturer’s recommendation unless stated otherwise. 293T cells were co-transfected with plasmids for MLV (pLRB302, 3 µg) or HIV-1 NL4-3 (2 µg) along with either 3 µg of pBJ5 MARCHs (mMARCH2; hMARCH2; mMarch2^N^–hMARCH2^C^; hMARCH2^N^–mMarch2^C^; mMARCH2 mutants: S18G, Q58P, C61G, N75C; hMARCH2 mutants: G18S, P58Q, G61C, C75N) or empty vector. To determine the effect of human MARCH2 polymorphisms, we co-transfected 293T cells with pNL4-3.Luc.R-E-(2.5 µg), pcDNA3.1 NL4-3 Env (1.25 µg) and either pBJ5 MARCH2 plasmids (1, 2, and 4 µg) or empty vector. Culture media was removed and replenished 24 h after transfection. Virus containing culture supernatants and cells were harvested 48 h after transfection. Culture supernatants were processed as described before (see *Virus preparation* section) and used for subsequent infection experiments. Cells were lysed in 1× RIPA buffer (150 mM NaCl, 1% NP-40, 0.5% sodium deoxycholate, 0.1% SDS, 25 mM Tris, pH 7.4, with Halt phosphatase and protease inhibitors). Cell lysates and virus pellets were mixed with 1 × sample loading buffer, boiled at 100 °C for 10 minutes and resolved on either 8%, 10%, 12%, or 15% sodium dodecyl sulfate-polyacrylamide gels (SDS-PAGE) as needed. Blots were probed using the following antibodies: goat anti-MLV gp70 (42), rat anti-MLV p15E (clone 42/114; Kerafast), rat anti-MLV p30 (R187, ATCC CRL-1912), rabbit anti-myc (Cell Signaling Technology), mouse anti-myc (Cell Signaling Technology), mouse anti-HIV gp41 (Chessie8) (NIH HIV Reagent program, ARP-526), human anti-HIV gp41 clone 2F5 (NIH HIV Reagent program, ARP-1475), mouse anti-HIV p24 (NIH HIV Reagent program, ARP-4121), rabbit anti-MARCH2 (Invitrogen, PA5-30620), rabbit anti-GAPDH 14C10 (Cell Signaling Technology), and monoclonal anti-β-actin (Sigma-Aldrich). Horseradish peroxidase (HRP)-conjugated anti-rabbit IgG (Cell Signaling Technology), HRP-conjugated anti-rat IgG (Cell Signaling Technology), HRP-conjugated anti-human IgG (Sigma-Aldrich), HRP-conjugated anti-mouse IgG (EMD Millipore), HRP-conjugated anti-goat IgG (Sigma-Aldrich), and light chain specific HRP-conjugated anti-mouse IgG (Cell Signaling Technology). Bands were detected using the enhanced chemiluminescence detection kits Clarity and Clarity Max ECL (Bio-Rad).

To determine the importance of the human MARCH2 domains on its antiviral function, we co-transfected 293T cells with a plasmid for HIV-1 NL4-3 (1.2 µg) along with pcDNA3.1 hMARCHs (250 ng of hMARCH2, hMARCH2 Δ2-30, hMARCH2 Δ31-56, hMARCH2 PDZ mutant; 80 ng of hMARCH2 RING-CH^mut^; 200 ng of hMARCH2 TM chimeras). For experiments with human MARCH2 isoforms, we co-transfected 293T cells with pNL4-3.Luc.R-E-(2.5 µg), pcDNA3.1 NL4-3 Env (1.25 µg) along with 700 ng of either pcDNA3.1 hMARCH2 isoform 1, pcDNA3.1 hMARCH2 isoform 2 or empty vector. In our experiments with HIV-1 envelope TM domain swap, we co-transfected 293T cells using 3 µg of either pcDNA3.1 NL4-3 Env or NL4-3 Env TM^TR^ along with either 2 µg of pBJ5 hMARCH2 or empty vector. Culture supernatants and cells were harvested 48 h post transfection and processed as described above.

### Luciferase assay

293T cells were co-transfected with plasmids for MLV molecular clone (pLRB302 3 µg), pFB-luc (1 µg), and 4 µg of either pBJ5 hMARCH2, mMARCH2 or pBJ5 mMARCH2 carrying the human amino acids at the indicated positions (S18G, Q58P, C61G, and N75C) or empty vector. Culture supernatants were harvested 48 h post transfection and processed as described above (see *Virus preparation* section). NIH 3T3 cells were seeded in 12-well plates (0.9 × 10^5^ cells/well). Cells were infected the next day with culture supernatants and lysed 48 hpi followed by measuring luminescence using the Steady-Glo luciferase assay system (Promega) per the manufacturer’s recommendation and a Biostack4 (BioTek) luminometer. Luminescence values were normalized to MLV p30^CA^ levels.

We generated pseudoviruses for experiments with human MARCH2 amino acids substitutions from that of mouse MARCH2 or mouse MARCH2 C61G by co-transfecting 293T cells with pNL4-3.Luc.R-E-(2.5 µg), pcDNA3.1 NL4-3 Env (1.25 µg) along with 4 µg of either pBJ5 mMARCH2, hMARCH2, hMARCH2 variants with the mouse MARCH2 amino acids at the indicated positions (G18S, P58Q, G61C, and C75N), mMARCH2 C61G or empty vector. Transfection conditions for pseudovirus production with human MARCH2 polymorphic variants and isoforms have been described above (see *Transfection and Immunoblotting section*). Culture supernatants were processed as described above and used to infect U373-MAGI-CXCR4 cells (0.5 × 10^5^ cells/well in a 24-well plate). Cells were lysed 48 hpi followed by measuring luminescence as described above. Luminescence values were normalized to HIV-1 p24^CA^ levels. MLV p30^CA^ or HIV p24^CA^ levels on culture supernatants was determined by immunoblotting (see *Transfection and Immunoblotting* section) followed by densitometry using ImageJ software (National Institutes of Health).

### Coimmunoprecipitations (coIPs)

For our coIP experiments examining endogenous human MARCH2, we pelleted 10 × 10^6^ H9 cells and infected with an 0.1 MOI of HIV-1 NL4-3 in a total volume of 2 ml complete RPMI media containing 2 µg of polybrene for 2 h at 37 °C. Cells were washed once to remove viral inoculum and maintained in 20 ml of complete RPMI media. Cells were harvested at 5 dpi, washed twice with cold 1× PBS and lysed in 1 ml of 1× RIPA buffer (150 mM NaCl, 1% NP-40, 0.5% sodium deoxycholate, 0.1% SDS, 25 mM Tris, pH 7.4, with Halt phosphatase and protease inhibitors). Following centrifugation, clarified lysates (1000 µg) were used for immunoprecipitation. CoIPs were performed using the Dynabeads protein A immunoprecipitation kit (Thermo Fisher Scientific) per manufacturer’s instructions with some modifications. Briefly, cell lysates were incubated with mouse anti-HIV-1 gp41 (Chessie 8) (NIH HIV Reagent program, ARP-526; 1: 200 dilution) overnight at 4 °C. Next day, 50 µl protein A Dynabeads was added and incubated for 1 h at RT. Dynabeads were then washed and eluted. The eluted fractions were subjected to western blot analysis. Antibodies used to probe western blots are described above (see *Transfection and Immunoblotting* section).

For our coimmunoprecipitation experiments examining the role of the human MARCH2 TM domains, we co-transfected 293T cells with 1.2 µg of HIV-1 molecular clone (pNL4-3) along with pcDNA3.1 hMARCH2 plasmids (100 ng of hMARCH2, hMARCH2 TM2^TR^; 25 ng of hMARCH2 TM1^M4^) or empty vector. At 48 h post-transfection, cells were lysed in IP lysis buffer (25 mM Tris pH 7.4, 150 mM NaCl, 2 mM MgCl_2_, 1% Triton X-100, 5% Glycerol, Benzonase [25 U/ml], 1× Halt protease inhibitor cocktail). Following centrifugation, clarified lysates (1000 µg) were incubated with 25 µl of Protein A Dynabeads for 6 h at 4 °C. Precleared lysates were incubated with mouse anti-HIV-1 gp41 (Chessie 8) (NIH HIV Reagent program, ARP-526; 1: 200 dilution) overnight at 4 °C. Next day, 30 µl protein A Dynabeads was added and incubated for 1 h at RT followed by 2 h incubation at 4 °C. Finally, Dynabeads were then processed as described above.

For coIPs examining the role of the HIV-1 TM domain, we co-transfected 293T cells with 4 µg of pcDNA3.1 NL4-3 Env, 3 µg of pcDNA3.1 NL4-3 Env TM^TR^, and 50 ng of pcDNA3.1 hMARCH2 or empty vector. Cells were harvested, precleared, and incubated with mouse anti-HIV-1 gp41 (Chessie 8) (NIH HIV Reagent program, ARP-526; 1: 200 dilution) as described above. Next day, 30 µl protein A Dynabeads was added and incubated for 2 h at RT followed by immunoblot analysis as described above.

### Cell surface staining and FACS analyses

Cell surface expression levels of NL4-3 Env and NL4-3 Env TM^TR^ chimera were performed as described previously with some modifications (24). Briefly, 293T cells (2.5 × 10^6^ /well) were seeded in a 12 well plate. Next day, cells were co-transfected with plasmids for NL4-3.Luc.R-E-(1 µg) and either NL4-3 Env (3 µg), NL4-3 Env TM^TR^ (2 µg) or empty vector. Cells were harvested 48 h post transfection, washed, incubated with 20 µg/ml of monoclonal anti-HIV-1 gp120 antibody VRC01 (NIH HIV Reagent Program, NIAID, NIH, ARP-12033) for 1 h at 4 °C. Cells were washed 3 times with fluorescence-activated cell sorting (FACS) buffer (1× PBS containing 2% FBS) followed by incubation with Alexa Flour 633 goat anti-human IgG (1: 2000 dilution, Invitrogen). Finally, cells were washed, fixed in 2% paraformaldehyde for 10 min at 4 °C, and acquired on BD FACSCelesta followed by data analysis using FlowJo version 10.8.0.

### PCR analyses for human MARCH2 isoforms

To detect isoforms of human MARCH2, PCR was performed using cDNAs from H9 and THP-1 cells and the following primers: hM2 isoform 1 (5’-CTCACAGAGTGGCTGAAGG-3’/ 5’-CGTCCAGAGGACATAGATGG-3’, hM2 isoform 2 (5’-CTCACAGAGGTCTCCTTCC-3’/5’-GTCTCCTCTGCCACCTTC-3’). Primers for GAPDH were also used as controls (see *HIV-1 infection, interferon treatment and human MARCH2 expression analysis* section).

### Statistical analyses

Statistical analyses were performed using GraphPad Prism software version 10.0. The statistical tests used to determine significance are described in the figure legends. A difference was considered to be significant for *P* values of ≤ 0.05.

## Acknowledgements

We thank Victoria Truong, Wade Sigurdson, Corey Knowles, Carlos Batista, Namrata Deka for technical support and Xinqi Liu for kindly providing us with reagents used for this project. Research was supported by National Institutes of Health grants R01 AI165161 (SS) and R56 AI165161 (SS), SUNY and University at Buffalo startup funds (SS). S.U. was supported by Research Grant for New Scholar (Grant No. RGNS 65–026) by the Office of the Permanent Secretary, Ministry of Higher Education, Science, Research and Innovation (OPS MHESI), Thailand Science Research and Innovation (TSRI) and Chulalongkorn University.

---

**Fig S1.**
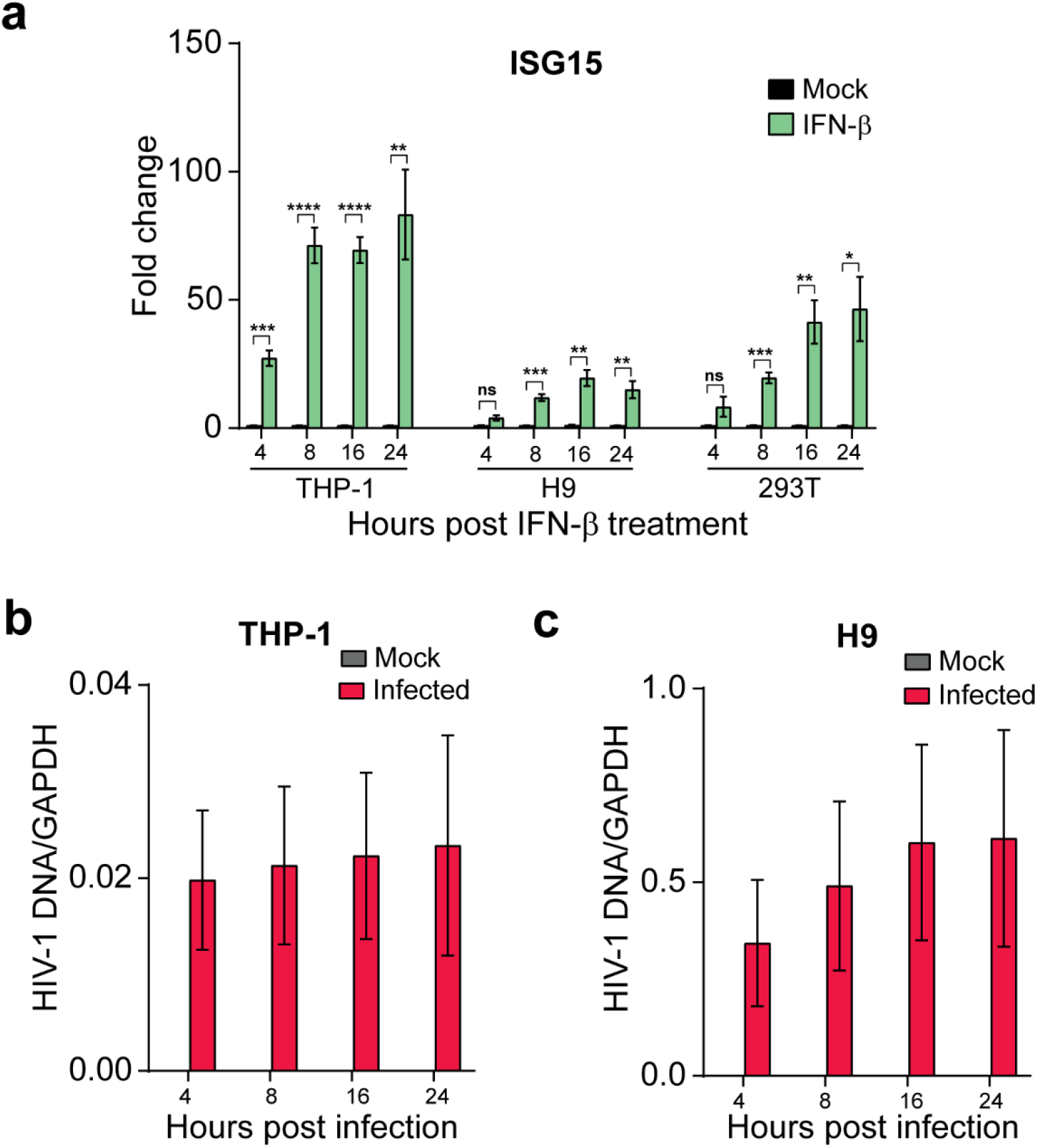
Confirmation of type I IFN response and HIV-1 infection in THP-1 and H9 cells. **(a)** Expression fold change of human *ISG15* transcript relative to mock, normalized to GAPDH in PMA-differentiated THP-1, H9, and 293T cells treated with human IFN-β (500 units/ml) for 4 h, 8 h, 16 h, and 24 h. **(b)** HIV-1^JR-CSF^ *nef* DNA relative to GAPDH in PMA-differentiated THP-1 cells at 4 h, 8 h, 16 h, and 24 h post infection. **(c)** HIV-1^NL4-3^ *nef* DNA relative to GAPDH in PMA-differentiated THP-1 cells at 4 h, 8 h, 16 h, and 24 h post infection. Mock indicates mock-treated (PBS). All graphs represent mean ± SEM from 3 independent experiments. Statistical analysis performed using unpaired t-test (two-tailed). ns, non-significant; *, *P* ≤ 0.05; **, *P* ≤ 0.01; ***, *P* ≤ 0.001; ****, *P* ≤ 0.0001.

**Fig S2.**
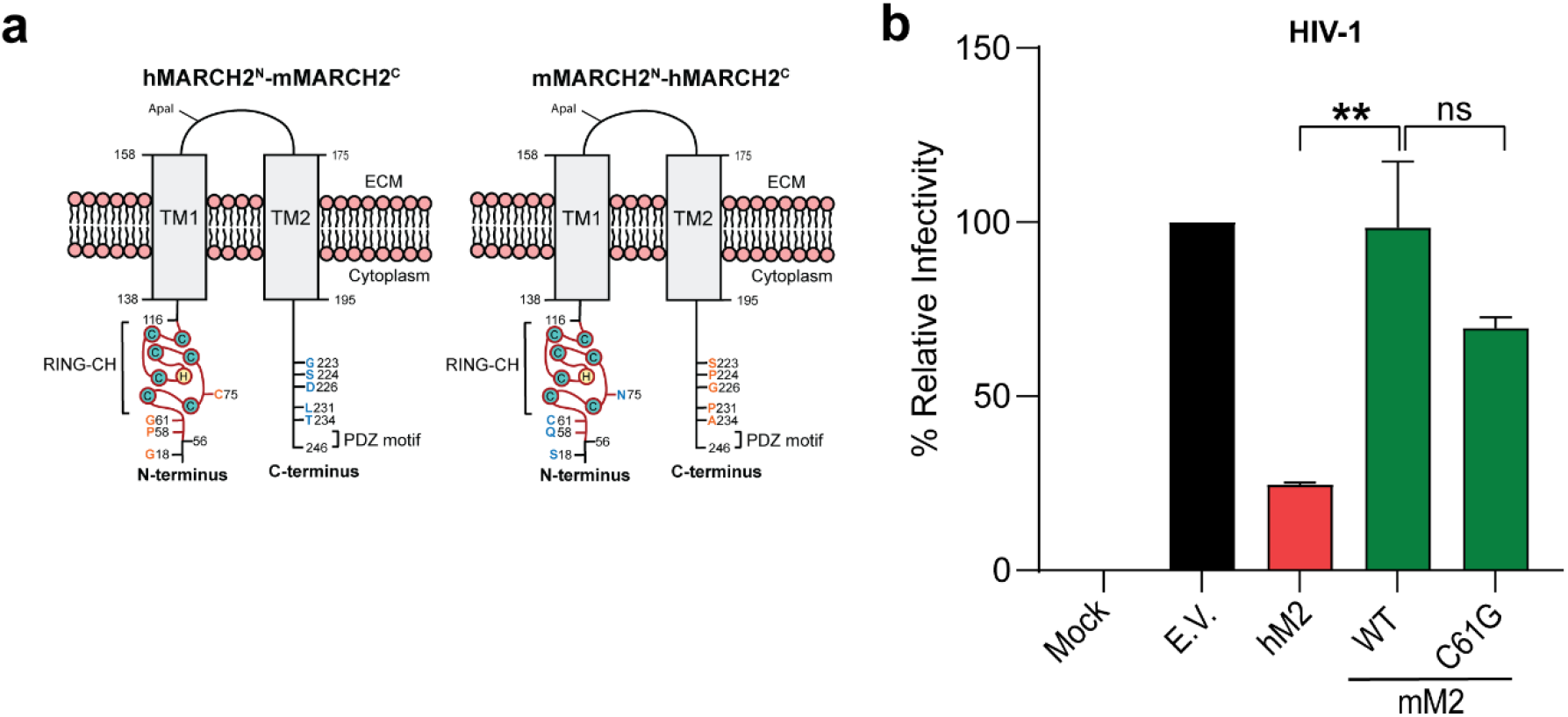
Mouse MARCH2 with C61G substitution has no anti-HIV-1 function. **(a)** Schematic diagram depicting the MARCH2 chimeras (hMARCH2^N^-mMARCH2^C^ and mMARCH2^N^-hMARCH2^C^) generated by swapping the N- and C-terminal cytoplasmic tails between human and mouse MARCH2 proteins. The positions of the differing amino acid of interest in human (orange) and mouse (blue) MARCH2 are indicated. **(b)** NL4-3 Env pseudotyped luciferase reporter viruses were generated in the presence of either wildtype human and mouse MARCH2, mouse MARCH2 C61G mutant or empty vector (E.V.). Pseudoviruses were used to infect U373-MAGI-CXCR4 cells, luciferase levels were measured 48 hpi and normalized to HIV-1 p24^CA^ levels of the input virus. The percentage (%) of relative infectivity was determined with respect to virus produced in the presence of E.V. Graphs represent means ± SD from 3 independent experiments. Statistical significance was determined by unpaired t-test (two-tailed). ns, non-significant; **, *P* ≤ 0.01. (Really Interesting New Gene-CH domain, RING-CH; Transmembrane domain, TM; Extra cellular matrix, ECM; PDZ binding motif, PDZ motif; human MARCH2, hM2; mouse MARCH2, mM2; wildtype, WT)

**Fig S3.**
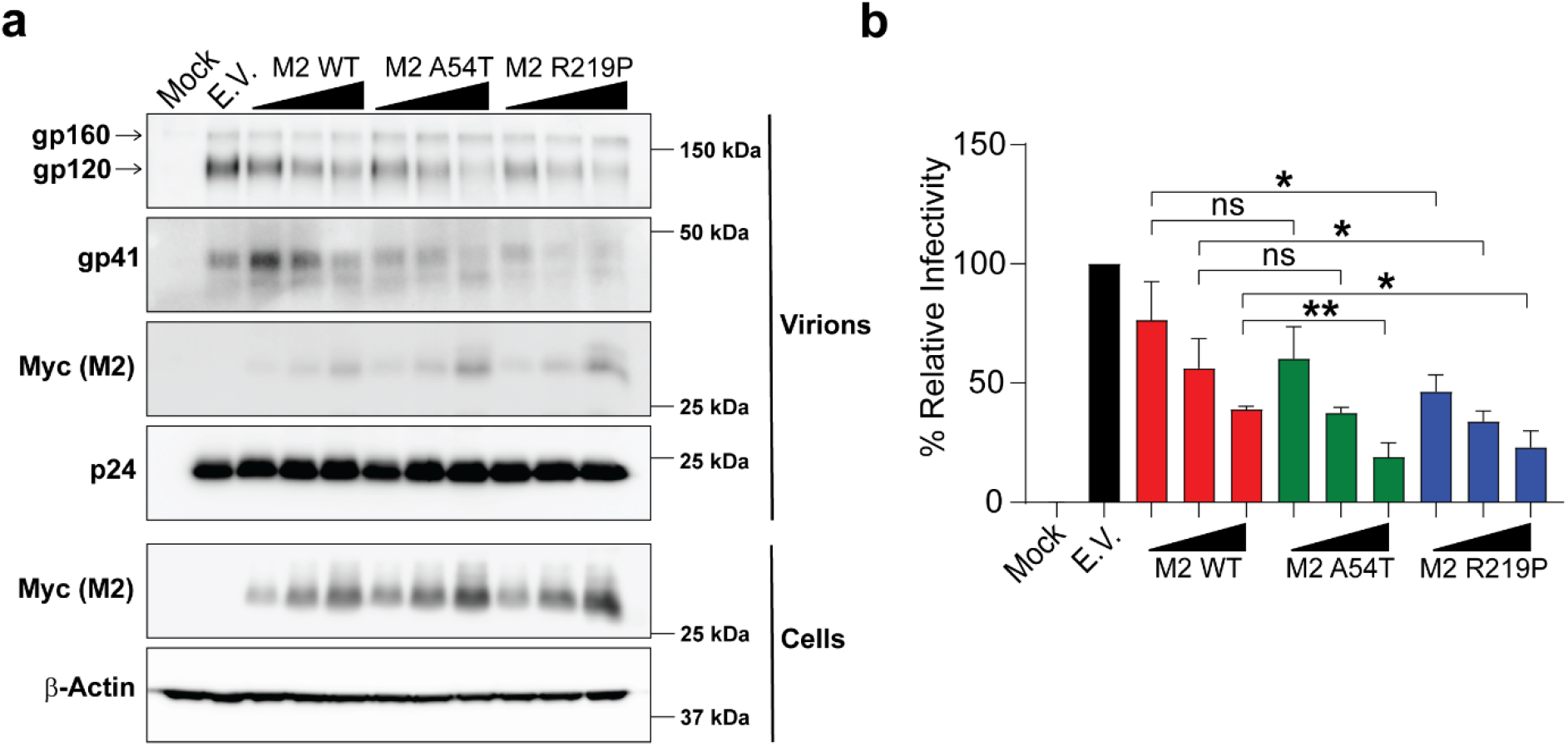
Naturally occurring MARCH2 polymorphisms restrict HIV-1 envelope. HIV-1 Env pseudoviruses were generated in the presence of either wildtype human MARCH2 (M2 WT), human MARCH2 polymorphic variants (M2 A54T, M2 R219P) or empty vector (E.V.). At 48 h post transfection, cells and culture supernatants containing pseudoviruses were harvested. **(a)** Indicated proteins in purified virion fractions and cell lysates were analyzed by immunoblotting. **(b)** Pseudoviruses were used to infect U373-MAGI-CXCR4 cells, luciferase levels were measured 48 hpi and normalized to HIV-1 p24^CA^ levels of the input virus. The percentage (%) of relative infectivity was determined with respect to virus produced in the presence of E.V. In **a**, representative gel images from 3 independent experiments are shown. In **b**, graphs represent means ± SD from three independent experiments. Statistical significance was determined by unpaired t-test (two-tailed). ns, non-significant; *, *P* ≤ 0.05; **, *P* ≤ 0.01.

**Fig S4.**
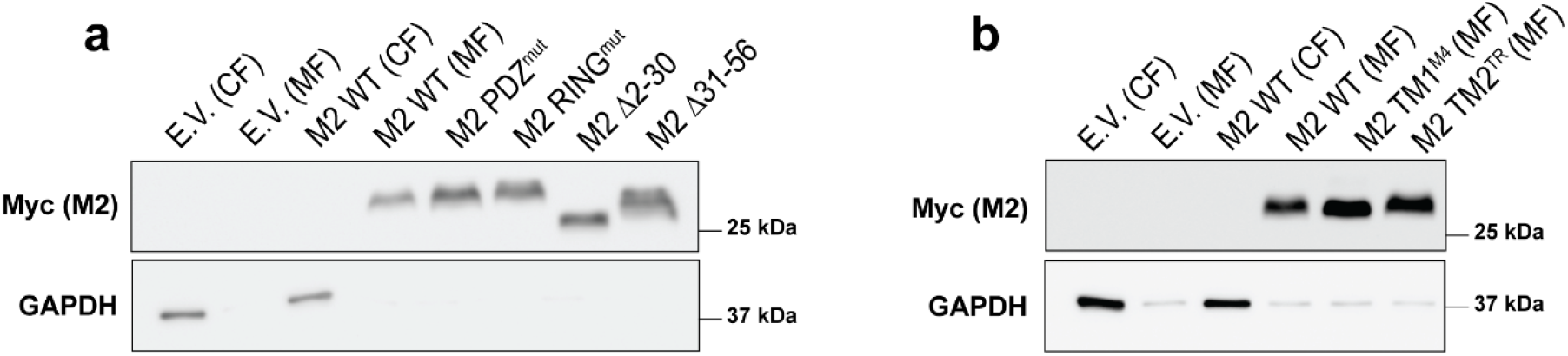
MARCH2 mutants localize to cellular membranes. 293T cells were transfected with either empty vector (E.V.), wildtype human MARCH2 (M2 WT), or the various MARCH2 mutants shown in Figure 3. Cells were harvested 48 h post transfection and integral membrane proteins were extracted using the MEM-PER Plus membrane extraction kit (Thermo Scientific) followed by immunoblotting. Membrane fraction purity was verified by probing for GAPDH. In both **a** and **b**, representative gel images from 2 independent experiments are shown. (membrane fraction, MF; cellular fraction, CF)

**Fig S5.**
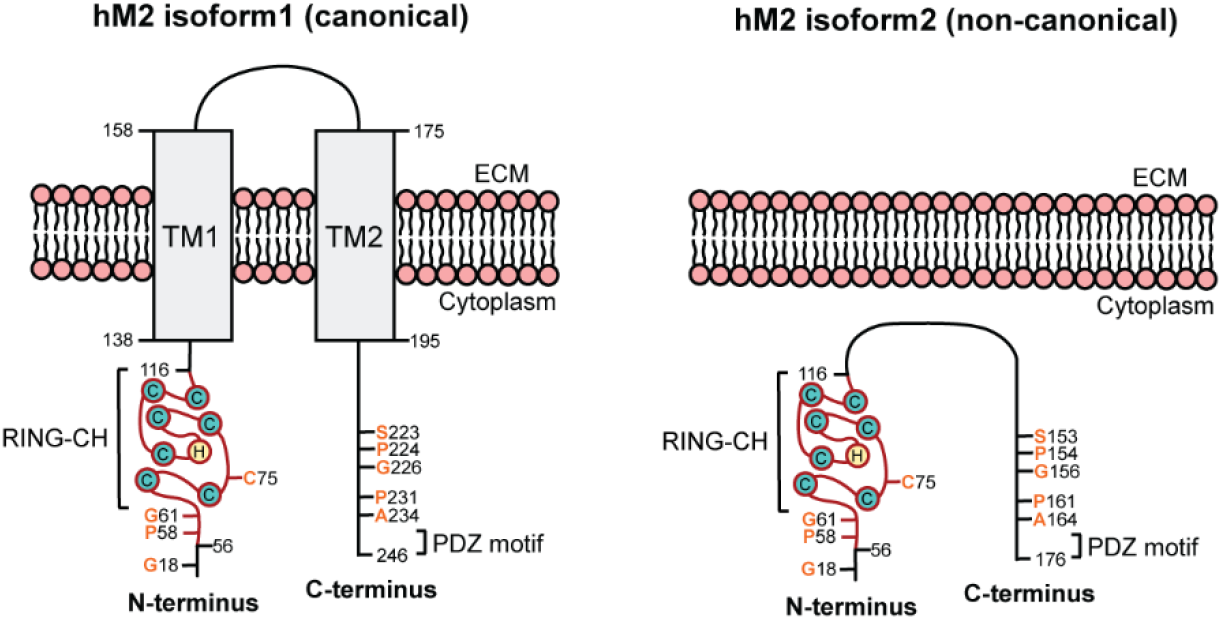
Isoforms of human MARCH2. Schematic diagram for human MARCH2 isoforms. The left panel represents the canonical (long) isoform 1 of human MARCH2, which consists of 246 amino acid residues and has two transmembrane domains (TM1 and TM2). The right panel represents the non-canonical (short) isoform 2 of human MARCH2 which lacks both TM domains (Δ125-194 residues from canonical isoform) resulting in a shorter polypeptide (176 amino acids residues).

**Fig S6.**
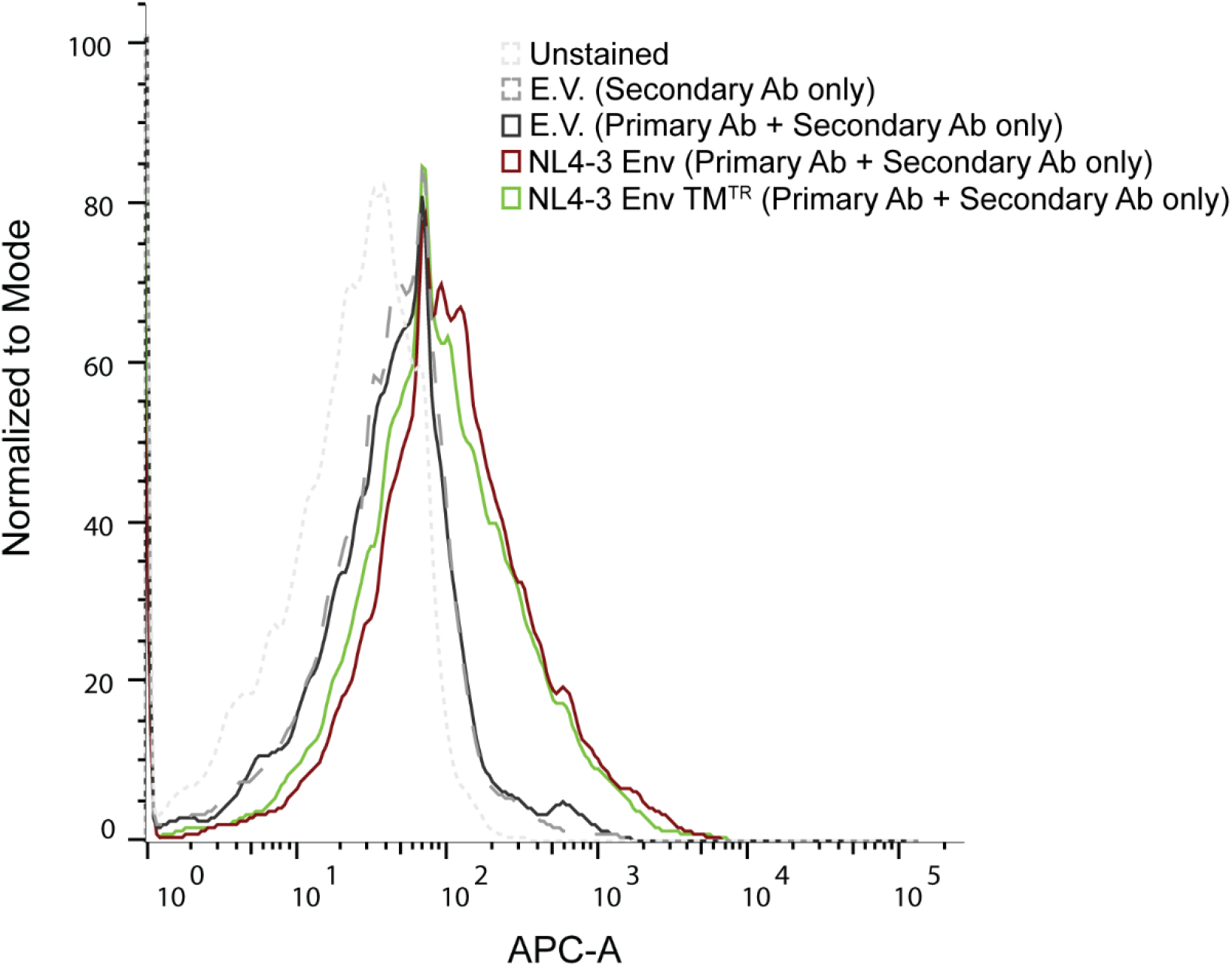
NL4-3 envelope mutant, in which the gp41 TM domain is replaced with that of human TR (NL4-3 Env TM^TR^) localizes at the plasma membrane. 293T cells co-transfected with plasmids for either NL4-3 Env, NL4-3 Env TM^TR^or empty vector (E.V.). Cells were harvested 48 h post transfection, washed, incubated with monoclonal anti-HIV-1 gp120 antibody VRC01 (primary Ab). Cells were stained with Alexa Flour 633 goat anti-human IgG (Secondary Ab) and then subjected to FACS. Shown are representative histograms from 3 independent experiments.

**Table S1:** The SNP frequencies of *MARCH2* at A54T (rs1133893) and R219P (rs34099346) obtained from 1000Genomes and ranked by alternative allele frequencies of each population (release version: 20230706150541)

| SNVs | Population | Frequencies |  | Sample Size |
| --- | --- | --- | --- | --- |
|  |  | Ref Allele | Alt Allele |  |
| NP_001356706.1:p.Ala54Thr<br>(rs1133893) | Global | G=0.8011 | A=0.1989 | 5008 |
|  | European | G=0.7078 | A=0.2922 | 1006 |
|  | American | G=0.728 | A=0.272 | 694 |
|  | East Asian | G=0.7619 | A=0.2381 | 1008 |
|  | South Asian | G=0.768 | A=0.232 | 978 |
|  | Africa | G=0.9652 | A=0.0348 | 1322 |
| NP_001356706.1:p.Ag219Pro<br>(rs34099346) | Global | G=0.9065 | C=0.0935 | 5008 |
|  | South Asian | G=0.826 | C=0.174 | 978 |
|  | American | G=0.833 | C=0.167 | 694 |
|  | European | G=0.8330 | C=0.167 | 1006 |
|  | East Asian | G=0.9980 | C=0.0020 | 1008 |
|  | Africa | G=0.9909 | C=0.0091 | 1322 |

**Table S2:**
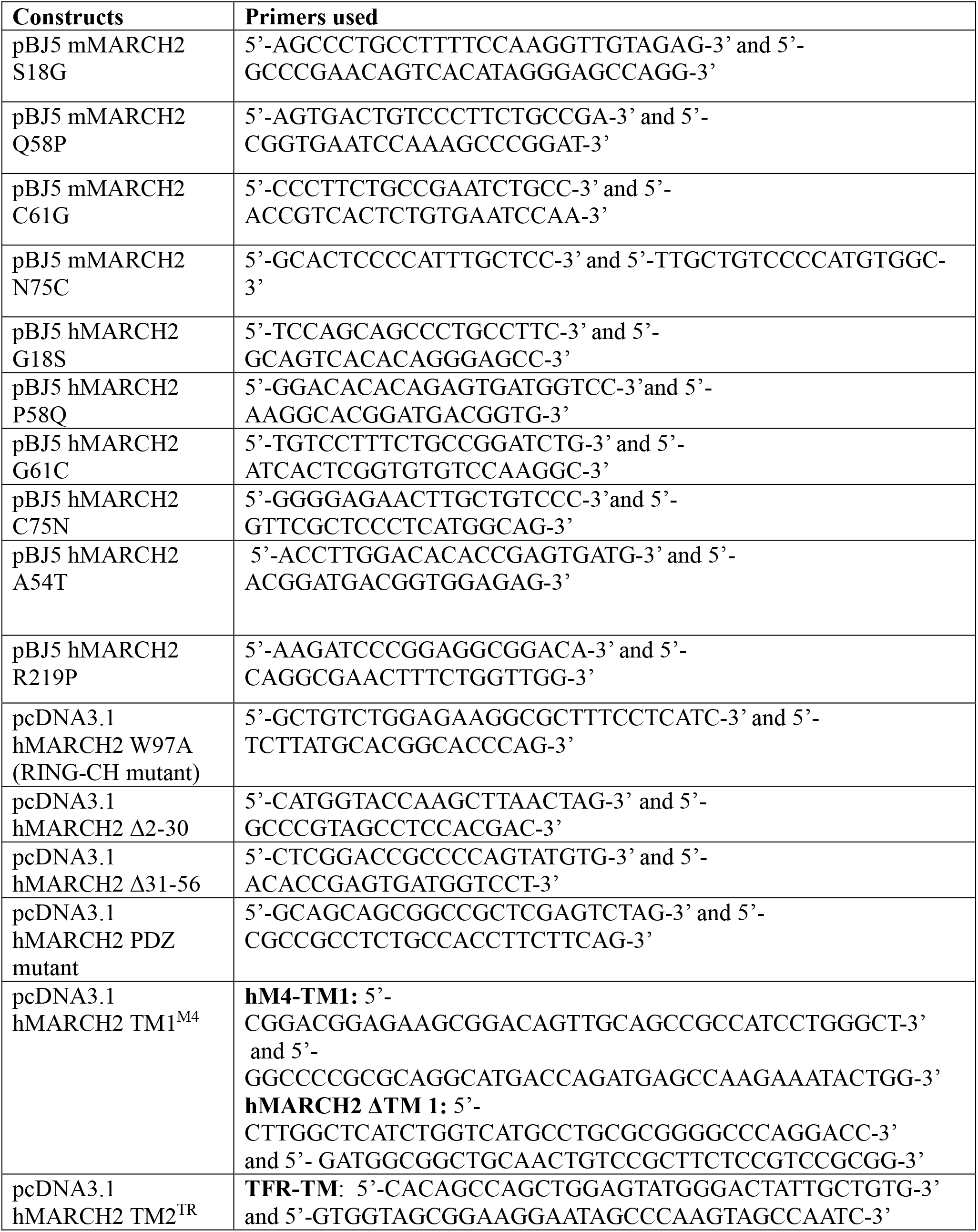
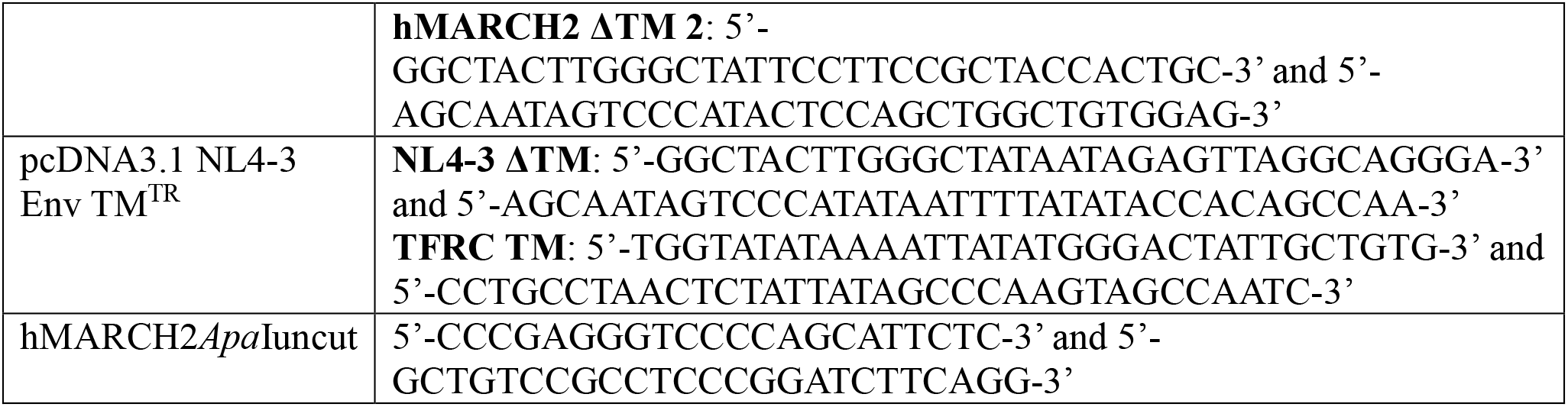
Primers used for generating MARCH2 variants.

## Notes

### Competing Interest Statement

The authors have declared no competing interest.

